# Stochastic Force Generation in an Isometric Binary Mechanical System: A Thermodynamic Model of Muscle Force Generation

**DOI:** 10.1101/2023.09.20.558702

**Authors:** Vidya Murthy, Josh E. Baker

**Affiliations:** University of Nevada, Reno School of Medicine, Department of Pharmacology

## Abstract

With implications for both clean energy technologies and human health, models of muscle contraction provide insights into the inner workings of one of the most energy-efficient engines on the planet and into the modifications to that engine that lead to human diseases. However, only scientific methods can provide these insights. A binary mechanical model is a recently developed thermodynamic model of muscle contraction that implies a novel entropic kinetic formalism, provides a solution to a paradox that has perplexed scientists for over a century, and accounts for many mechanical and energetic aspects of muscle contraction. Here we use this model to perform discrete state chemical simulations of isometric force generation under different conditions and show explicitly that force generating kinetics are bounded by thermodynamic equations, that four phases of force generation occur as four separate thermodynamic processes, and that periodic force generation emerges with amplitudes and periodicities that bifurcate between constant and stochastic values through mechanisms easily understood relative to ideal thermodynamic processes. We discuss these results relative to experimental observations of spontaneous oscillatory contractions (SPOCs) in muscle and periodic force generation in small myosin ensembles.

**Significance Statement:** Most models of muscle contraction to date are based on the obsolete 17th century scientific philosophy that the force of the system is determined by the force of the molecules in that system. A new thermodynamic model of muscle provides a completely different interpretation of muscle mechanics and chemistry, implies a novel thermodynamic kinetic formalism, and has solved a paradigm that has intrigued scientists for over a century. Here, we use this model to simulate muscle force generation and show that force generating kinetics are constrained by thermodynamic equations that provide a clear mechanism for the periodic force generation that emerges from these stochastic simulations.

## INTRODUCTION

Muscle is a complex and dynamic macromolecular system that is integral to physiological functions such locomotion, digestion, and cardiac contractility. The molecular mechanism for muscle contraction is well established through single molecule mechanic (1–4) and kinetic (5–7) studies. Specifically, strong actin binding to myosin gated by inorganic phosphate, P, release (MDP to AMD) induces a discrete conformation change (a large and distinct rotation of the myosin lever arm) in myosin that displaces elements external to the motor a distance of 8 nm (8–11). If the external element displaced is elastic, the motor generates force proportional to the stiffness of that external elastic element (11). The question remains, how is this simple two-state binding mechanism (Fig. 1) related to the chemistry and mechanics of muscle contraction?

**Figure 1.**
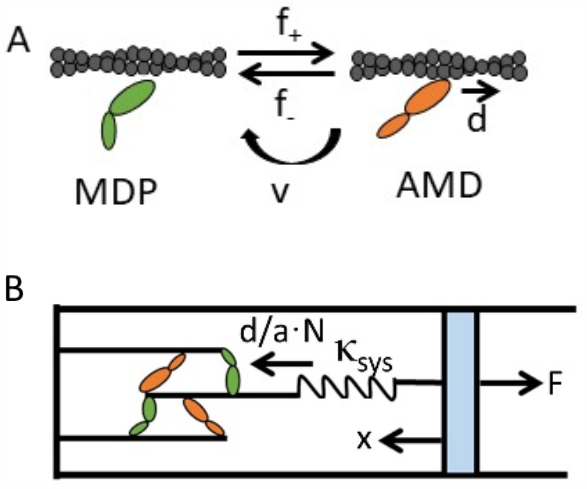
A binary mechanical model. (A) Two-state (MDP and AMD) scheme in which actin (gray helix) binding to a single myosin motor (ovals) induces a myosin lever arm rotation that displaces elements external to that myosin a distance, *d*. The reversible binding reaction occurs with forward, *f*_*+*_, and reverse, *f*_*–*_, rates. Myosin motors irreversibly detach from actin through an active (ATP-dependent) process at a rate *v*. (B) Muscle force is represented by a single spring with stiffness κ_sys_ that on one end (left) is reversibly stretched a distance *d*/(*aN*) with each binding step while the other end is defined by macroscopic mechanics (e.g., held at a fixed force or length).

In 1938, A.V. Hill proposed a thermodynamic model of muscle contraction (12). Challenging A.V. Hill’s model, in 1957, A.F. Huxley proposed a molecular mechanic model of muscle contraction that violates the principles of thermodynamics (13), which is why in 1974 T.L. Hill was compelled to formalize a new kind of molecular (not ensemble) mechanochemistry(14). While historically A.V. Hill’s model is based on Gibbs’ chemical thermodynamics, Huxley’s model is based on a corpuscular mechanic philosophy proposed by Boyle in the 17^th^ century (15) that assumes the mechanical properties of a system are determined by the mechanical properties of the molecules within that system. Although corpuscular mechanics has been widely dismissed as an obsolete scientific idea, a corpuscular theory of muscle contraction continues to be defended (16, 17) and provides the formal foundation for most models of muscle contraction to date (17).

An ensemble of molecular mechanical switches constitutes a binary mechanical system (18) – a model that while analogous to a quantum spin system has only recently been developed. This model accounts for most chemical and mechanical aspects of muscle contraction such as the steady state force-velocity relationship, work loops (e.g., cardiac pressure-volume loops), and the four phases of a force transient following a rapid mechanical or chemical perturbation of muscle (18). The model also has broad physicochemical implications, implying a novel entropic kinetic formalism (18). This reformulation has provided the first explicit solution to the Gibbs (mixing) paradox (19).

Gibbs described thermodynamics as the mechanical laws of a macroscopic system of particles as measured by an observer, not the mechanical laws of particles as conjured by corpuscularians. We have shown that the laws of mechanics of muscle can be described by a single system spring with a force, *F* (Fig. 1B) (18). One side of the spring (Fig. 1B, right) describes the macroscopic state (force and length) of muscle, while the other side of the spring (Fig. 1B, left) is collectively stretched by binding steps of individual motors. When muscle is held at a fixed length (right side of spring), molecular force generation (left side of spring) occurs adiabatically and is described by a simple linear force equation (force increases linearly with the number of bound motors) (18). When the muscle system is at equilibrium, the system force (right side of the spring) is described by the Gibbs binding reaction free energy. Under one or the other of these idealized conditions, muscle force generation follows one or the other ideal thermodynamic pathway (Eqs. 1 and 2). Under non-ideal conditions, muscle force generation follows intermediate pathways.

Mathematical and continuous models have been developed (8, 18) that describe smooth trajectories but do not capture the emergent stochastic mechanics of myosin motor ensembles. Therefore, using a binary mechanical model here we develop discrete chemical simulations of isometric muscle force generation. We perform these simulations over a wide range of conditions and observe that force generation follows four different thermodynamic pathways (adiabatic, isothermal, ergodic, and catastrophic) where each pathway corresponds to a different phase of force generation. This level of complexity is remarkable considering that all four processes emerge from a single binding mechanism (a molecular switch).

Under certain conditions, we observe periodic force generation like that observed in small myosin motor ensembles (20, 21) and with spontaneous oscillatory contractions (SPOCs) in muscle (22). The periodicities and amplitudes of simulated periodic force generation bifurcate between constant and stochastic values with mechanisms that become evident when we compare stochastic simulations to ideal thermodynamic processes.

The analysis below is a theoretical exploration of the force-generating thermodynamic processes that naturally emerge from a simple binary mechanical model. None of these processes can be described with a corpuscular mechanic analysis. As such, this analysis establishes a fundamentally new theoretical toolbox for modeling force generation in muscle and molecular motor ensembles. While the goal here is to characterize the foundational processes of a new theory, in the discussion we compare our simulations with observations of force generation in muscle and small myosin motor ensembles. We show that the model reconciles disparate force-generating behaviors recently observed in two different in vitro force studies.

## METHODS

We assume the two-state kinetic scheme illustrated in Fig. 1A. The reversible formation of a strong bond between actin (A) and myosin (M) induces a conformational change (i.e., a myosin lever arm rotation) in myosin that displaces elements external to the motor a distance, *d*, of 8 nm (8, 9, 11, 23). If the external elements are elastic, this binding step (the working step) reversibly generates force. The reversible working step (actin-myosin binding) occurs with ADP (D) bound to myosin and is gated by the release of inorganic phosphate (P) (4, 9) with forward, *f*_*+*_, and reverse rates, *f*_*–*_. Through the ATPase reaction cycle, irreversible detachment of actin and myosin occurs with ADP release and ATP induced actin-myosin dissociation at an effective rate *v*. In these simulations, we assume ATP hydrolysis is not rate limiting.

In thermodynamics, equations of state describe relatively simple macroscopic relationships between the force, *F*, temperature, pressure, and chemical energetics of a system of molecules. In a thermodynamic model of muscle contraction, a single spring describes muscle force, *F*(18). Figure 1B illustrates a single spring of stiffness κ_sys_ that at one end (right side) is defined by the macroscopic mechanical state of the system (force or fixed length) while the other end (left side) is collectively stretched by reversible working step displacements.

When muscle (the right side, Fig. 1B) is held at a fixed length (isometric), working steps collectively generate force through sequential displacements, *d*/(*aN*), of the system spring (left side, Fig. 1B) where *N* is the number of myosin motors and *a* is the system ergodicity(18). Ergodicity, *a*, describes the extent to which within a given chemical state {*N*_*MDP*_,*N*_*AMD*_} all microstates (i.e., the number of ways that out of *N* myosin motors *N*_*AMD*_ are strongly bound to actin) are equally probable. An ergodic equilibrium (*a* = 1) requires that the system force is dynamically equilibrated among all *N* myosin motors at which point the effective step size is *d*/*N*. At a minimum, the system force is generated by one of *N* myosin motors, in which case the effective step size is that of one motor, *d*, and the ergodicity, *a*, has a minimum value of 1/*N*. Thus, ergodicity, *a*, in a mechanical system can be viewed as the fraction of *N* motors over which the system force is dynamically equilibrated.

Two ideal thermodynamic force-generating relationships can be defined: adiabatic and isothermal. Isometric (or adiabatic) force increases linearly with the number, *N*_*AMD*_, of bound myosin motors as

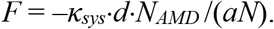

In our simulations, both *F* and *N*_*AMD*_ are initialized to 0. In the current analysis, we assume that force, *F*, varies linearly with ergodicity, *a*, as *F* = *aF*_*o*_ (i.e., the more myosin motors over which force is dynamically equilibrated, the higher the force), where *F*_*o*_ is the ergodic equilibrium force (when *a* = 1). Adiabatic force is then

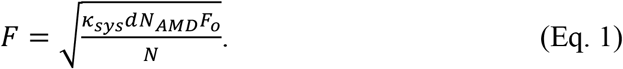

This ideal, adiabatic force curve is plotted (substituting *N*_*AMD*_ = *N* – *N*_*MDP*_) in Fig. 2A (blue line).

**Figure 2.**
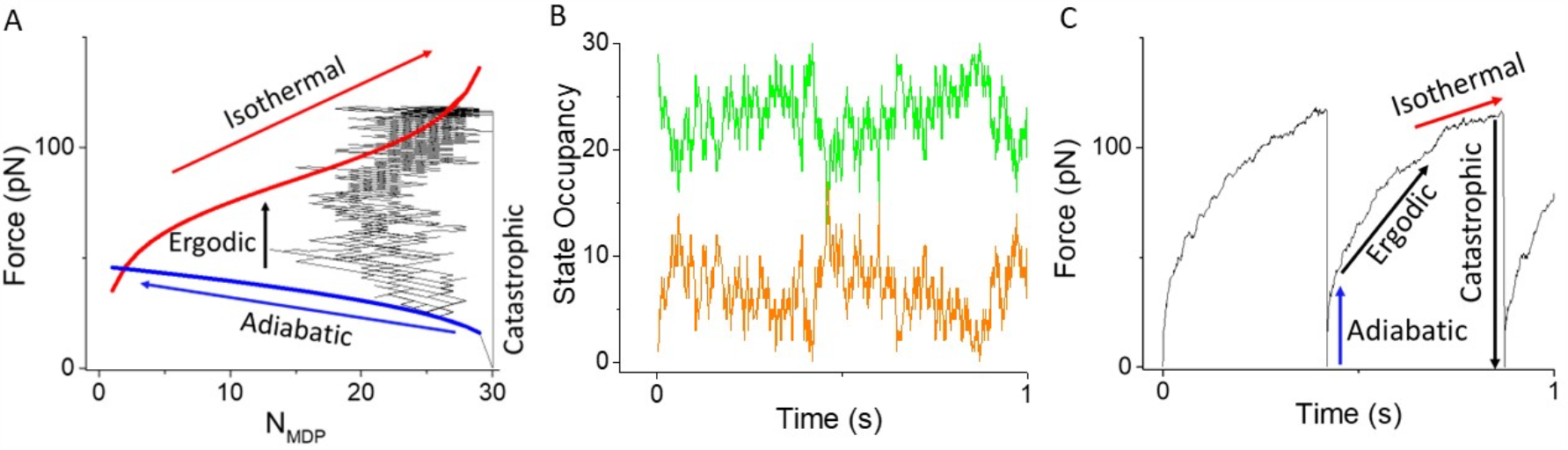
Simulations of force-generating loops overlaid with Eqs. 1 and 2. (A) Plots of Eqs. 1 (blue curve) and 2 (red curve) with *N* = 30, κ_sys_ = 2 pN nm^-1^, *a* = 0.5 are overlaid with replots (black trace) of *N*_*MDP*_ and F from Panels B and C. (B) *N*_*MDP*_ as a function of time from simulations of the model in Fig. 1 using parameters in panel A and *v* = 50 s^−1^. (C) Force plotted as a function of time from the simulation in panel B.

The reaction free energy for binding (Fig. 1A) is(18)

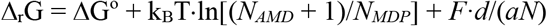

where ΔG° is the standard reaction free energy and *N* = *N*_*MDP*_ + *N*_*AMD*_, defining an equilibrium (Δ_r_G = 0) force

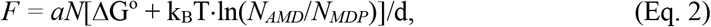

where here *N*_*AMD*_ + 1 is approximated as *N*_*AMD*_. When the system force is dynamically distributed among all myosin motors (*a* = 1) *F* reaches an ergodic equilibrium binding isotherm (Fig. 2A, red line) at a force *F* = *F*_*o*_. When force generation equilibrates (Δ_r_G = 0) before an ergodic state is reached (a < 1), *F* stalls at a force *F* = *aF*_*o*_ (a non-ergodic equilibrium).

Equations 1 and 2 describe two different relationships between force, *F*, and chemistry, *N*_*AMD*_, defined for two different ideal conditions (adiabatic or isothermal). Physiological muscle function, however, is non-equilibrium and often non-ideal(18), which is to say it is bracketed by but does not follow ideal pathways (Eqs. 1 and 2). While these physiological processes are not defined by analytical expressions (Eqs. 1 and 2) they can be described computationally through kinetic simulations of the mechanochemical steps defined by Eq. 1 with kinetic rates defined by Eq. 2.

The ratio of the forward, *f*_*+*_, and reverse, *f*_*–*_, working step rates is

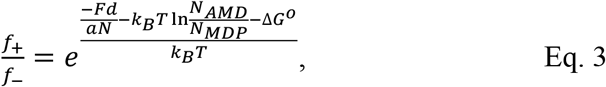

where Eq. 3 is the probability of the muscle system being found in state {*N*_*MDP*_–1,*N*_*AMD*_+1} relative to state {*N*_*MDP*_,*N*_*AMD*_} along a system reaction energy landscape tilted by the energetic components of the reaction free energy. According to.

Eq. 3, adiabatic force generation stalls when *f*_*+*_ = *f*_*–*_ at a non-ergodic force (substituting *F* from Eq. 2 into Eq. 3).

Discrete simulations are based on the two-state mechanochemical model in Fig. 1, using rate constants defined by Eq. 3. Specifically, Eq. 3 can be rewritten

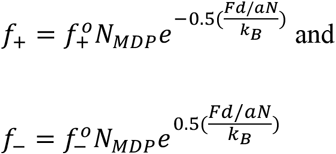

where *f*_*+*_^*o*^ and *f*_*–*_^*o*^ are forward and reverse rate constants. The parameter *a* is set to *F*/*F*_*o*_ and limited to a minimum value of 1/*N* and a maximum value of 1. In all simulations, we use model parameters consistent with experimental studies: *f*_*+*_^*o*^ = 30 s^−1^ and *f*_*–*_^*o*^ = 0.1 s^−1^ (rates consistent with a ΔG° of –5.7 k_B_T) and *d* = 8 nm (1, 24).

## RESULTS

From simulations of a two-state binding reaction (Fig. 1), we plot time traces of *F* (Fig. 2B) and the occupancies of the AMD (*N*_*AMD*_, orange line) and MDP (*N*_*MDP*_, green line) states (Fig. 2C). The data in Figs. 2B and 2C are then replotted as *F* versus *N*_*MDP*_ in Fig. 2A, showing that force generation initially follows the ideal adiabatic pathway (Eq. 1), transitions from adiabatic to isothermal pathways along a non-ideal ergodic pathway, and then generates force along the isothermal pathway before a catastrophic loss of force occurs when the last bound myosin head (at *N*_*MDP*_ = 1) detaches. Below we dissect the components of this force generating loop and the conditions under which it occurs. We begin by simulating isometric force generation through the binding reaction alone (*v* = 0), assessing mechanisms and key factors that affect them.

In Fig. 3, we consider the effects of the system spring stiffness, *κ*_*sys*_, on the binding reaction. The left panels in Fig. 3 are simulated time courses of both force, *F*, and the equilibration parameter, *a*, (inset), and the right panels are *F* vs. *N*_*MDP*_ from the same simulations overlaid with ideal thermodynamic processes (Eq. 1 and 2). Panels from top to bottom are simulations performed at *κ*_*sys*_ ranging from 0.01 to 10 pN/nm.

**Figure 3.**
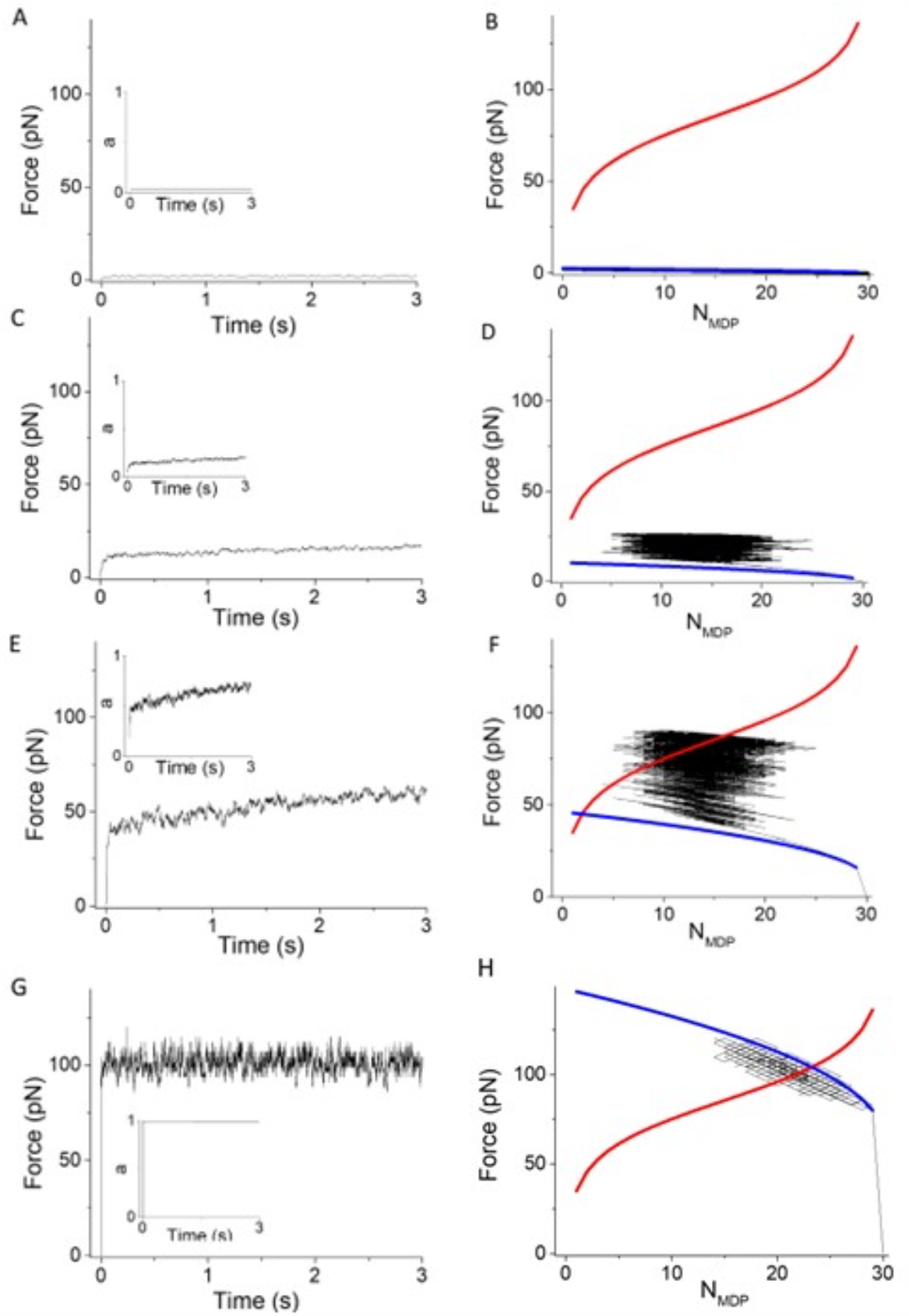
The effects of system spring stiffness, κ_sys_, on simulations of the model in Fig. 1, using *v* = 0, *N* = 30. Left panels are simulated force traces and the equilibration parameter, *a* (inset), and right panels *F* and *N*_*MDP*_ from the same simulations overlaid with Eqs. 1 (blue curve) and 2 (red curve) using κ_sys_ values of (A and B) 0.01 pN/nm, (C and D) 0.16 pN/nm, (E and F) 2 pN/nm, and (G and H) 10 pN/nm.

When the system spring stiffness is low (Figs. 3A and 3B, *κ*_*sys*_ = 0.01 pN/nm) myosin heads rapidly bind actin with no significant force generation and a value for *a* that never exceeds 1/*N* (*a* < 1/*N* or *F*/*F*_*o*_ > 1/*N* or *F* < *F*_*o*_/*N*). Because the total force, *F*, generated by *N* myosin motors never exceeds the equilibrium force of one myosin motor, *F*_*o*_/*N*, the maximum system force is unable to balance (i.e., stop) the chemical force of even one motor. Thus, the reaction proceeds to *N*_*AMD*_ = *N* as it would in solution in the absence of mechanical force.

At higher stiffnesses (Figs. 3C and 3D, *κ*_*sys*_ = 0.16 pN/nm), the value *a* rapidly exceeds 1/*N*, and adiabatic force generation stalls at a non-ergodic isotherm when *N*_*AMD*_ = *N*_*MDP*_. Here, the mechanical force is balanced – not against the equilibrium binding isotherm (Δ_r_G = 0, *a* = 1) but – against a non-ergodic force, *F* = *aF*_*o*_ (Δ_r_G = 0, *a* < 1). The mechanism is that a relatively large non-equilibrium effective step, *d*/(*aN*), against a relatively small non-equilibrium force, a*F*_*o*_, performs mechanical work, *aF*_*o*_*d*/(*aN*), equal to that performed at equilibrium, *F*_*o*_*d*/*N*, by a relatively small equilibrium effective step *d*/*N* against the equilibrium force, *F*_*o*_.

Here, force generation stalls after the system samples only one of the many ways in which half the myosin motors can bind actin. In other words, only one microstate of the chemical state {*N*_*MDP*_=*N*/2;*N*_*AMD*_=*N*/2} is accessible, which means the system has equilibrated (Δ_r_G = 0) in a non-ergodic state (*a* < 1). Absent an active process (Δ_r_G < 0), the system likely remains in this frustrated state because the likelihood that all bound myosin motors reversibly detach with equal probability, *f*_*–*_, is low (forces are not dynamically distributed among all motors). If this were possible, however, our simulations imply a mechanism for spontaneous ergodicity (driven by *a* < 1) through which *F* approaches *F*_*o*_ and *a* approaches 1. This occurs because the effective displacement, *d*/(*aN*), by a forward step is larger than that of a reverse step since the effective step size is larger (*a* is smaller) at lower forces. At sufficiently high stiffness, adiabatic force generation directly equilibrates with the equilibrium isotherm in an ergodic state (Figs. 3G and 3H).

In Fig. 4, we consider the effects of increasing the number, *N*, of myosin motors on simulated binding reactions. The left panels are plots of the simulated time course for both force, *F*, and the equilibration parameter, *a*, (inset), and the right panels are *F* vs. *N*_*MDP*_ from the same simulations overlaid with ideal thermodynamic processes (Eq. 1 and 2). Panels from top to bottom are simulations performed with *N* values ranging from 5 to 100 myosin motors. At low *N* (*N* = 5) adiabatic force generation directly equilibrates with the equilibrium isotherm with no ergodic transition required, whereas at higher *N* the ergodic force required to reach the equilibrium isotherm increases with *N*.

**Figure 4.**
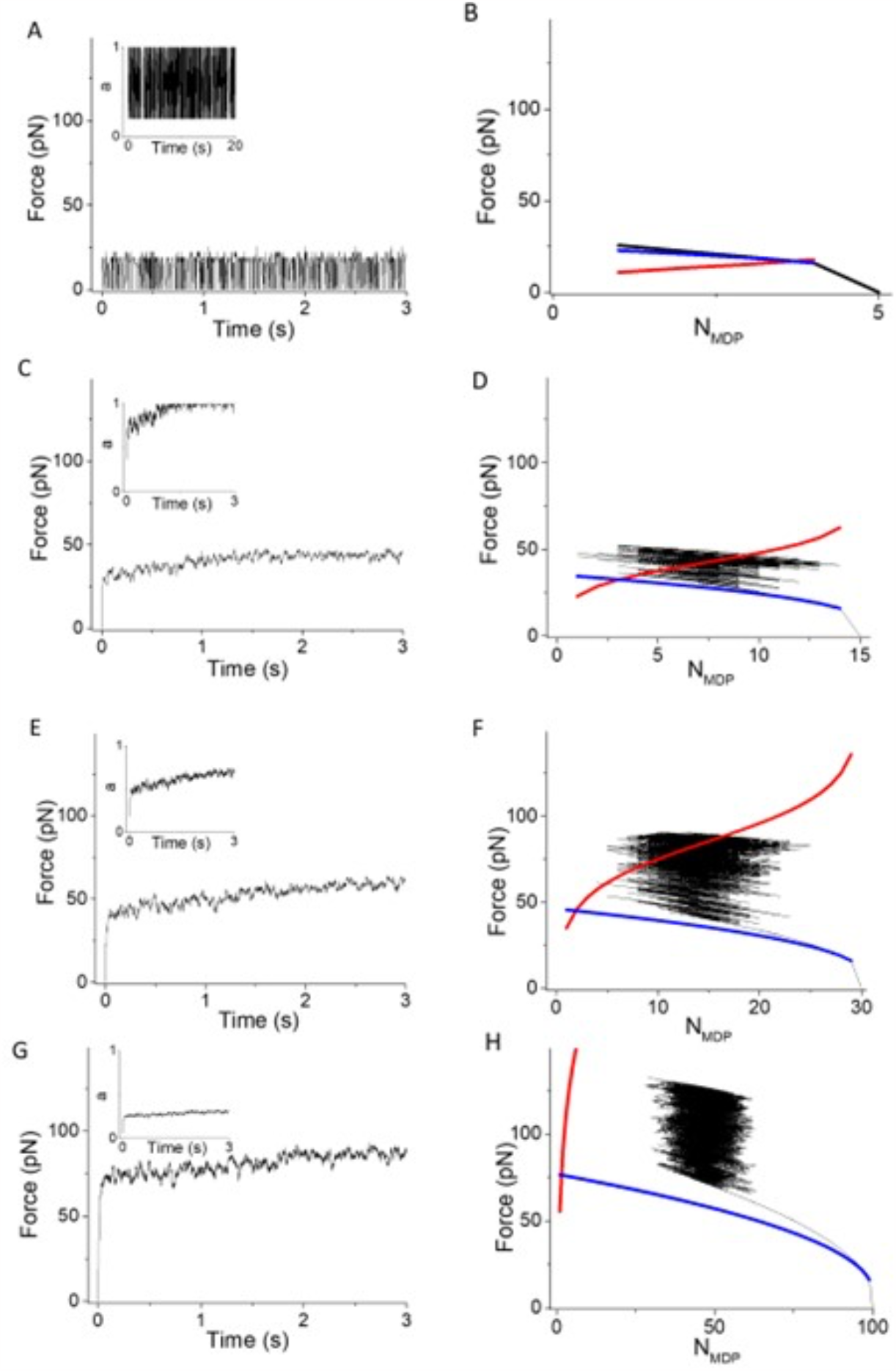
The effects of *N* on simulations of the model in Fig. 1, using *v* = 0, κ_sys_ = 2 pN/nm. Left panels are simulated force, *F*, and equilibration parameter, *a* (inset), and and right panels are *F* and *N*_*MDP*_ from the same simulations overlaid with Eqs. 1 (blue curve) and 2 (red curve) using *N* values of (A and B) 5, (C and D) 15, (E and F) 30, and (G and H) 100.

Figure 5 shows simulations of the model in Fig. 1 introducing irreversible rates, *v* (i.e., reaction cycles). The left panels are plots of the simulated time course of both force, *F*, and the equilibration parameter, *a*, (inset), and the right panels are *F* versus *N*_*MDP*_ from the same simulation overlaid with the ideal thermodynamic processes (Eqs. 1 and 2). Panels from top to bottom are simulations with *v* increasing from 1 to 100 sec^−1^.

**Figure 5.**
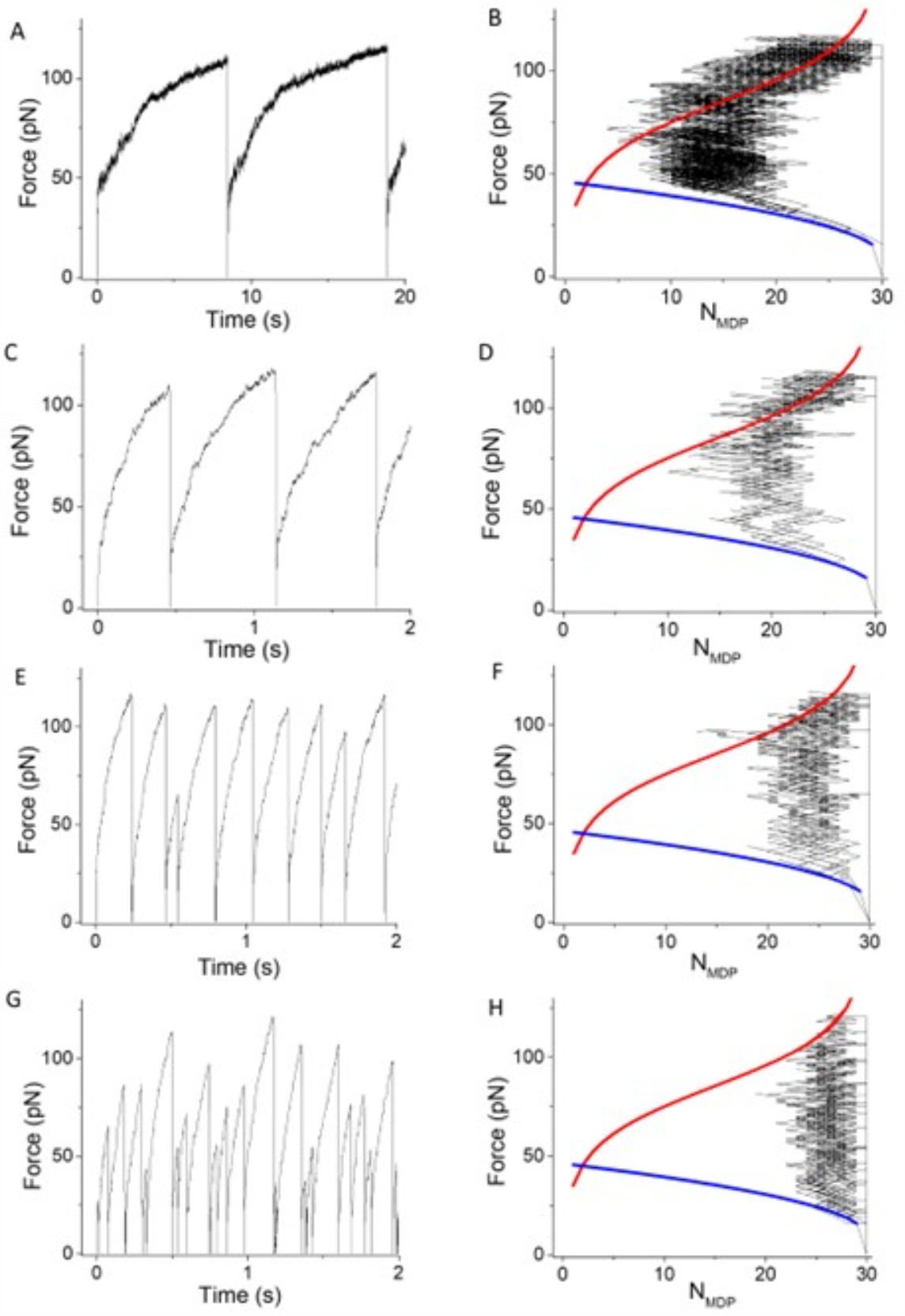
The effects of the irreversible detachment rate, *v*, on simulations of the model in Fig. 1, using *N* = 30, κ_sys_ = 2 pN/nm. Left panels are simulated force traces and the equilibration parameter, *a* (inset), and right panels are *F* and *N*_*MDP*_ from the same simulations overlaid with Eqs. 1 (blue curve) and 2 (red curve) using *v* equal to (A and B) 1 s^-1^, (C and D) 25 s^-1^, (E and F) 100 s^-1^, and (G and H) 200 s^-1^.

At low *v*, simulated force generation resembles that when *v* = 0 (Figs. 5E and 5F), occurring rapidly along the adiabatic curve until reaching the ergodic isotherm. In these simulations, free energy, Δ_r_G < 0, drives ergodicity by actively distributing forces through irreversible detachments. However, as described below, force generation need not reach an ergodic equilibrium through an active process, stalling instead at a non-ergodic force, *F* = *aF*_*o*_, in a frustrated state, *a* < 1. If an ergodic force, *F* = *F*_*o*_, is reached, irreversible detachments of myosin motors continue to perturb the binding reaction from equilibrium, and force continues to be generated along the binding isotherm, which according to Eq. 2 corresponds to a decrease in the number of bound myosin motors. Isothermal force generation continues until the last bound myosin motor detaches at which point a catastrophic loss of force, *F*, returns the system to its initial state. This four-stage force-generating loop repeats, resulting in periodic force generation. At low *v*, the amplitude of periodic force generation is the maximum force along the isotherm (Eq. 2, *N*_*MDP*_ = *N* – 1). The periodicity is limited by the rate, *v*, at which ergodic force and the maximum isothermal force are generated.

With increasing *v* (Fig. 5, top to bottom) irreversible detachment pulls the adiabatic binding reaction further from a non-ergodic equilibrium (*N*_*MDP*_ > *N*_*AMD*_ and Δ_r_G < 0) right shifting the force generating loop (right panels). That is, force generation along the adiabatic curve stalls at a larger *N*_*MDP*_; isentropic force generation occurs at a larger non-equilibrium *N*_*MDP*_; and less isothermal force generation is required to reach catastrophic force loss. Thus, with increasing *v* smaller loops and faster rates, *v*, decrease the period of force generating loops with little effect on the maximum amplitude since the force along the isotherm at which the last bound myosin motor detaches does not change with *v* (right panels).

At sufficiently high rates, *v*, isentropic force generation occurs at relatively large *N*_*MDP*_ values from which catastrophic force loss occurs stochastically (the probability that all myosin motors detach through a stochastic mechanism is comparable to the probability that all motors detach through an isothermal mechanism). In Figs. 5E and 5F (*v* = 100 sec^−1^) one out of nine force generating loops terminates through a stochastic mechanism, decreasing both the period and amplitude of that loop. In Figs. 5G and 5H (*v* = 200 sec^−1^) most force generating loops terminate stochastically and exhibit stochastic periods and amplitudes. This transition to stochastic periodic force generation occurs sooner (at a lower *v*) when *N* or *f*_*+*_ are lower or when *κ*_*sys*_ or *f*_*–*_ are higher.

Simulations in Figs. 6 and 7 illustrate the effects of stiffness, *κ*_*sys*_, and the number of myosin motors, *N*, on the amplitude and period of force generating loops. Figure 6 shows that increasing *κ*_*sys*_ decreases the period but not the amplitude of force generating loops because increasing *κ*_*sys*_ increases the slope of the adiabatic force generating curve (Eq. 1, Fig. 6 right panels blue curves) without affecting the isotherm (Eq. 2, Fig. 6 right panels red curves), creating smaller loops (shorter periods) of the same amplitude. Figure 7 shows that increasing *N* decreases the period and increases the amplitude of force generating loops because increasing *N* increases the isotherm (Eq. 2, Fig. 7 right panels red curves), creating larger loops (longer periods) with larger amplitudes.

**Figure 6.**
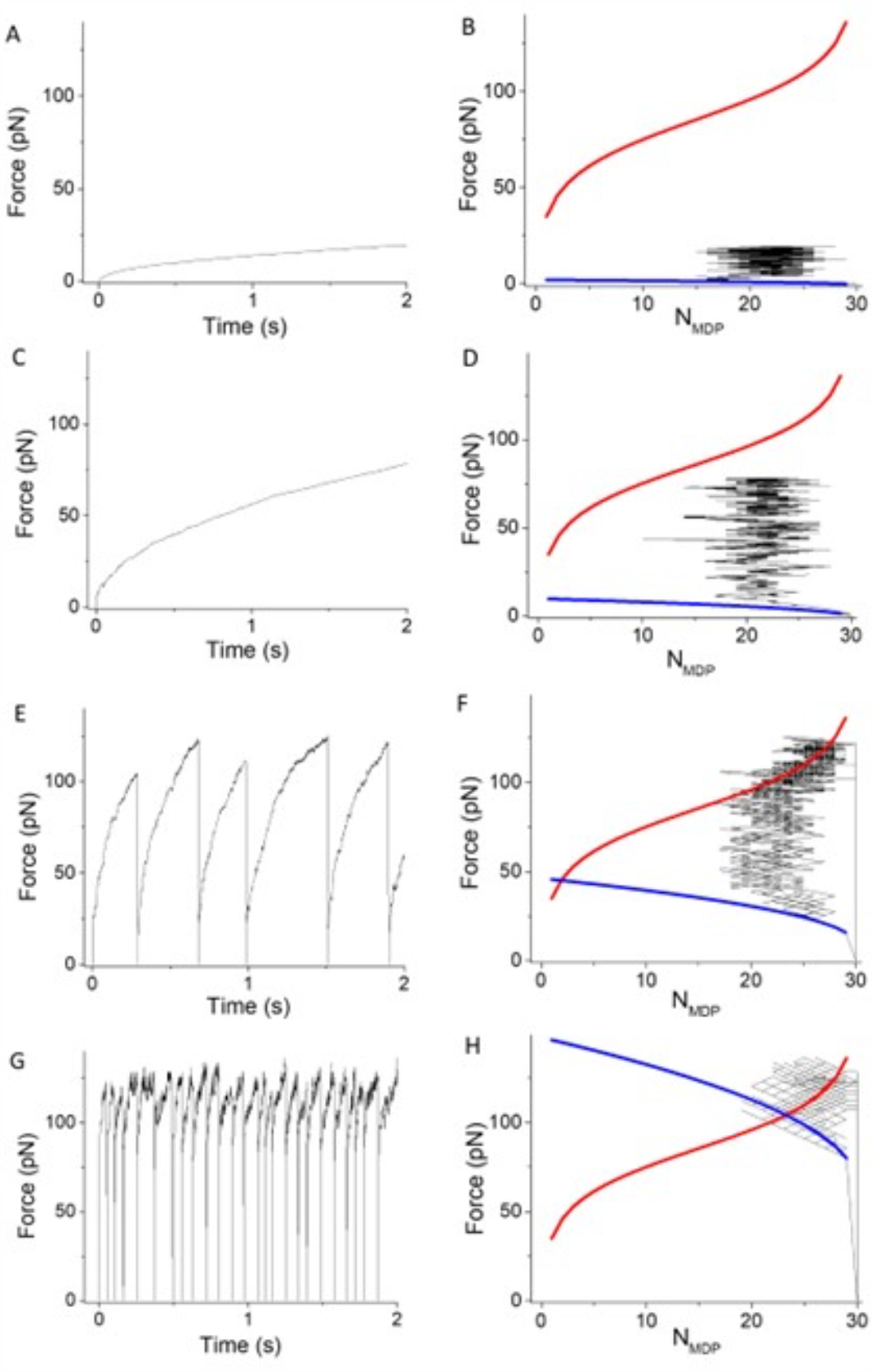
The effects of system spring stiffness, κ_sys_, on simulations based on the model in Fig. 1, using *v* = 50 s^-1^, *N* = 30. Left panels are simulated force traces and the equilibration parameter, *a* (inset), and right panels are plots of Eqs. 1 (blue curve) and 2 (red curve) using the parameters above overlaid with simulated parameters *F* and *N*_*MDP*_ at κ_sys_ values of (A and B) 0.01 pN/nm, (C and D) 0.16 pN/nm, (E and F) 2 pN/nm, and (G and H) 10 pN/nm.

**Figure 7.**
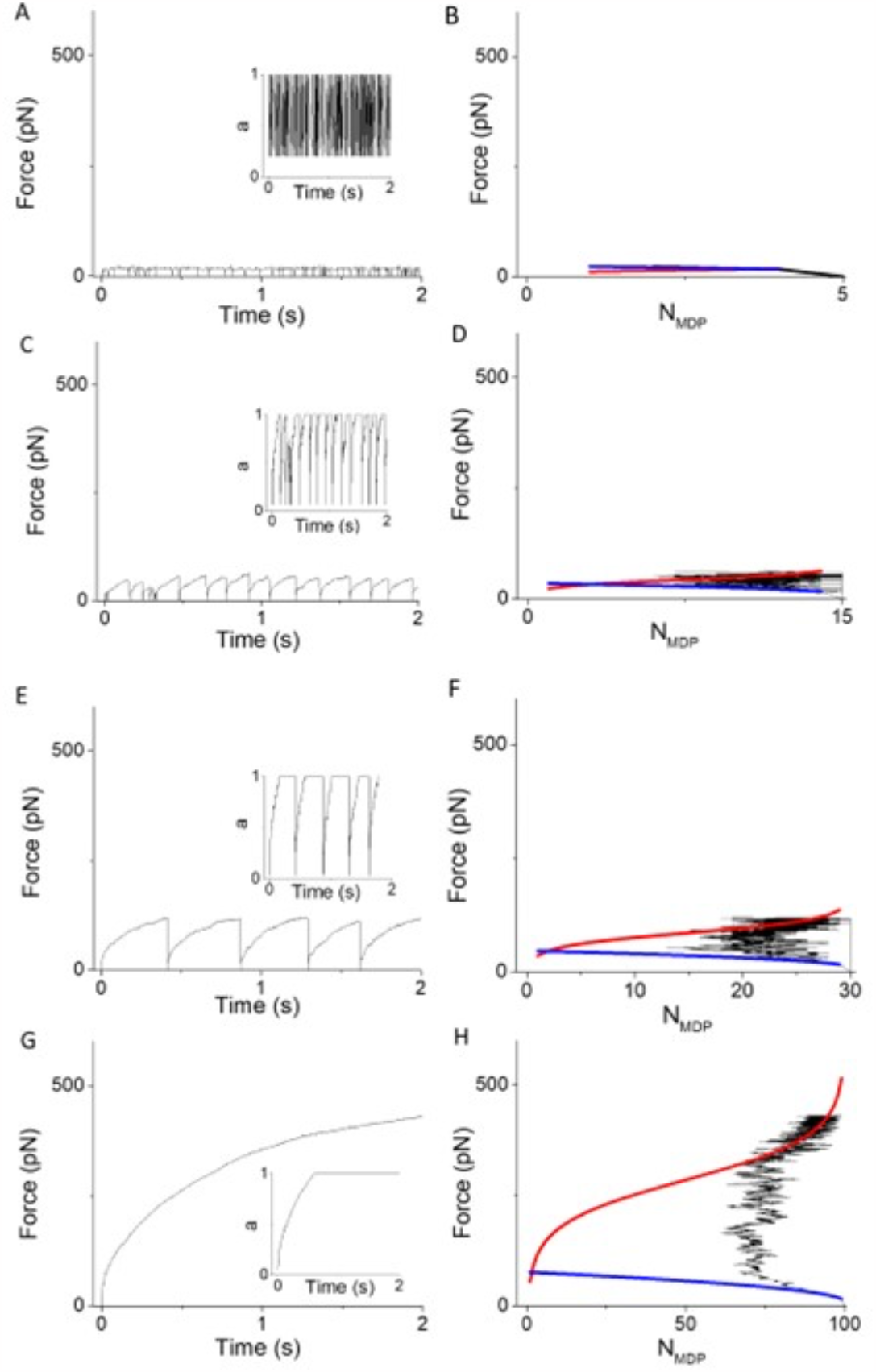
The effects of *N* on simulations based on the model in Fig. 1, using *v* = 50 s^-1^, κ_sys_ = 2 pN/nm. Left panels are simulations of force traces and corresponding equilibration parameter, *a* (inset), and right panels are plots of Eqs. 1 (blue curve) and 2 (red curve) using the parameters above overlaid with simulated parameters *F* and *N*_*MDP*_ at *N* values of (A and B) 5, (C and D) 15, (E and F) 30, and (G and H) 100.

## DISCUSSION

Here, we have presented discrete simulations of a reversible two-state binding reaction that displaces a system spring (Eq. 1) with forward and reverse rates derived from the binding free energy (Eq. 2). Four phases of force generation emerge from this single force-generating mechanism: adiabatic, isentropic, ergodic and catastrophic. Two of these phases are ideal thermodynamic processes. Adiabatic (isometric) force generation is described by the force equation (Eq. 1, Fig. 2 blue curve), and isothermal (equilibrium) force generation is described by the binding free energy (Eq. 2, Fig. 2 red curve). These ideal processes provide reference equations relative to which all other non-equilibrium, non-ideal force-generating processes are characterized.

In general, periodic force generation emerges from simulations of the two-state reaction cycle (Fig. 1A). Here we have explored the parameter space of periodicities and amplitudes and have determined mechanisms by comparing simulations to ideal processes. We observe that by changing certain parameters, periodicities and amplitudes shift between constant and stochastic values. The mechanism for this bifurcation is evident from a comparison of simulations to ideal processes.

In thinking about the physiological relevance of force generating loops, the beating heart often comes to mind. However, periodic force generation in muscle held at a fixed length is, in fact, the opposite isothermal process from that of the work loop of a heart. A work loop is a counterclockwise loop in Fig. 2A(18), whereas periodic force generation is a clockwise loop. In other words, the work performed by myosin motors on the system spring through force-generating loops is lost as heat through catastrophic force loss not as work on the surroundings through muscle shortening.

Periodic force generation has been observed in in vitro mechanics studies of small myosin motor ensembles(20, 21) and in spontaneous oscillatory contractions in muscle(22), and there are many fundamental similarities between our simulations and these observed behaviors. Broadly, if force generation occurred through a binding reaction alone (adiabatic force generation, Fig. 2A blue curve), the probability that all myosin motors spontaneously detach from actin would become exceedingly small at high forces with large numbers of myosin motors bound to actin. Clearly an isothermal phase is required through which high forces energetically reverse the force-generating binding reaction, consistent with Le Chatellier’s principle.

The in vitro mechanics studies of Kaya and colleagues show that small myosin motor ensembles generate periodic force generation that is both multiphasic and appears to bifurcate between constant and stochastic periodicities and amplitudes(20, 25). These experimental studies also demonstrate that the effective step size decreases with increasing force, consistent with our assumption of a force-dependent ergodicity, *a*. Our simulations provide clear, testable predictions for how experimental variables (e.g., optical trap stiffness, *κ*_*sys*_, the detachment rate, *v*, the number of myosin motors, *N*, etc) affect the phases, step sizes, periods, and amplitudes of periodic force generation measured in this assay.

A binary mechanical model accounts for the apparent discrepancy between the periodic force generation observed by Kaya and colleagues (20) and the constant non-ergodic stall force observed by Lombardi and colleagues (21) in a similar in vitro force assay. As described above, intramolecular forces prevent myosin motors from equilibrating with *F*_*o*_ (the force generated against other motors diminishes the force that can be generated in the system spring), stalling force generation in a non-ergodic (*a* < 1) state, *F* = *aF*_*o*_. In experiments of Kaya and colleagues, intramolecular forces were minimized by flexible S2 domain tethers, making the ergodic state accessible and allowing periodic force generation. In experiments of Lombardi and colleagues, there are no flexible tethers, and intramolecular forces stall force generation in a non-ergodic state, making the equilibrium isotherm inaccessible and preventing periodic force generation. We have shown that intramolecular interactions increase with decreasing [ATP], which accounts for Kaya’s observation of stalled non-ergodic forces at low [ATP].

Force is continually generated in active isometric muscle and must through some mechanism be lost as heat. If intramolecular forces prevent *a* from reaching 1, heat is dissipated with the disruption of intramolecular forces which is what energetically allows muscle force to stall at a constant *F* = *aF*_*o*_. If there are no intramolecular forces that prevent *a* from reaching 1, heat must be dissipated through the more catastrophic loss of force that occurs when all myosin motors are forced off actin through an isothermal process. In isometric muscle it remains unclear whether a constant isometric force results from a non-ergodic force or from asynchronous force-generating loops summed over many actin filaments.

The intramolecular forces that prevent ergodicity in isometric muscle resemble the intramolecular forces that prevent myosin motors from equilibrating (*a* = 0) with the external force, *F* = 0, in unloaded shortening muscle. The magnitude of this intramolecular force is *aF*_*o*_, where *a* was first defined by A.V. Hill as the coefficient of shortening heat (intramolecular forces are dissipated as heat during muscle shortening) with values ranging from 0.2 to 0.3 among different muscle types (24).

According to Eq. 3, at low *v*, adiabatic force generation stalls (*f*_*+*_ = *f*_*–*_) at *N*_*AMD*_/*N* = 0.5 in a non-ergodic state. That is, Δ_r_G reaches zero before *F* reaches the equilibrium (*a* = 1) force, *F*_*o*_. With an increase in the irreversible detachment rate, *v*, the overall rate of detachment increases, which drops *N*_*AMD*_/*N* below 0.5, and pulls the system further from equilibrium (Δ_r_G < 0), resulting in active ergodic force generation that is more effectively driven by Δ_r_G < 0 than by *a* < 1 (Fig. 5 vs. Figs. 2E and 2F). A positive correlation between *F, κ*_*sys*_, and *N*_*AMD*_ occurs only during adiabatic force generation. During ergodic force generation, F increases independent of *κ*_*sys*_ and *N*_*AMD*_, and during isothermal force generation, *F* increases with decreasing *N*_*AMD*_ independent of *κ*_*sys*_. The average value for *N*_*AMD*_/*N*, which is often erroneously referred to as a “duty ratio” (26), is primarily determined by the value for *N*_*AMD*_ during ergodic force generation.

There are several obvious discrepancies between our simulations and experimental observations. First, in in vitro force assays, relatively long periods with no mechanical activity are observed between force-generating loops. This is easily accounted for in simulations (27) by assuming a slower binding rate when all myosin motors are detached and actin filaments are no longer held in close proximity to the surface. With the goal of characterizing periodicity and amplitude, we did not include this conditional rate in our simulations. Second, in Figs. 6 and 7 we used a relatively slow rate, *v* (= 50 s^−1^), to study the effects of *N* and *κ*_*sys*_ on periodicity and amplitude. A more physiological *v* (= 200 s^−1^) at *N* = 5 results in the larger number of single binding events observed experimentally.

Individually, myosin motors function as mechanical switches that displace external elastic elements upon binding actin (9, 11, 18). Collectively, molecular mechanical switches function as a binary mechanical system (18). Despite being a logical extension of a quantum spin system, a binary mechanical model has only recently been developed. Characterizing the mechanochemical performance of this model system is, if nothing else, an intriguing exercise in physical chemistry. Demonstrating that it accurately accounts for muscle chemistry and mechanics makes it an important problem in biology and medicine. We have shown that a binary mechanical system accounts for the muscle force-velocity relationship, muscle force transients, muscle work loops, and here periodic (and non-periodic) muscle force generation (18). That these diverse and complex mechanochemical behaviors all emerge from a single molecular mechanism is remarkable. Because corpuscular mechanics cannot describe these thermodynamic processes, this simple binary mechanical model uniquely describes the thermodynamics of muscle and collective myosin motor function.

## Notes

### Competing Interest Statement

The authors have declared no competing interest.

